# CN-RNN: a Deep Learning Framework for Copy Number Variation Detection with Exome Sequencing Data

**DOI:** 10.64898/2026.05.13.724920

**Authors:** Dayuan Wang, Fei Qin, Wenhan Bao, Rhonda Bacher, Dongjun Chung, Qing Lu, Philip A. Efron, Guoshuai Cai, Feifei Xiao

## Abstract

Copy number variations (CNVs) are major structural genomic variants that contribute to a wide range of human diseases. Accurate detection of CNVs from whole-exome sequencing (WES) data has been a long-sought goal for clinical and population genetic studies. Despite recent progress, existing WES-based CNV callers still suffer from high false-positive rates and reduced recall for short-length variants, and current deep learning methods have not fully used complementary information in region-level genomic features. Here we present CN-RNN, a deep learning-based CNV caller for WES data. The model combines a bidirectional long short-term memory (BiLSTM) branch that captures local depth changes and contextual dependencies across neighboring exons with a parallel multi-layer perceptron (MLP) branch that encodes region-level metadata such as GC content, mappability, and exon length. CN-RNN was trained on the Autism Sequencing Consortium (ASC) parent-child trio cohort using the Mendelian rule of inheritance to ensure high-quality training sets. It was evaluated across three independent datasets, in which we showed that CN-RNN outperformed existing WES-based CNV callers and deep learning methods. CN-RNN offers a scalable, accurate tool for CNV profiling in WES-based studies and supports broader application of CNV analysis in population and clinical research. CN-RNN is available at https://github.com/FeifeiXiao-lab/CN-RNN.

## Introduction

Copy number variants (CNVs), which involve duplications or deletions of DNA segments in the genome, are a significant component of human genetic variation. These variants comprise up to 10% of the human genome [1]. They can significantly impact gene expression, phenotypic diversity, and susceptibility to diseases, including various cancers [2, 3], congenital heart defects [4], neurodevelopmental disorders [5] and schizophrenia [6]. Accurate profiling of CNVs is essential for understanding the genetic architecture of complex diseases and advancing precision medicine.

Whole-exome sequencing (WES) has been widely employed in large-scale disease studies and clinical diagnostics due to its cost-effectiveness and direct functional interpretation. WES captures protein-coding exons, however our ability to accurately detect CNVs is limited due to sparse and discontinuous coverage caused by high variability in the read depths driven by GC content effects, batch effects, and mappability differences [7–9]. Numerous computational tools such as CORRseq [10], EXCAVATOR2 [11], XHMM [12], CoNIFER [13], and CODEX2 [14] have been developed to address these challenges. Still, due to the high dimensionality, technical biases, and low signal-to-noise ratio in WES data, existing methods often generate inconsistent calls and excessive false positives.

Meanwhile, a growing number of machine learning and deep learning (DL)-based approaches have been proposed for improved CNV detection. For example, CN-Learn integrates outputs from multiple CNV callers using a random forest classifier and improves precision by leveraging genomic features among callers [15]. More recently, convolutional neural networks (CNNs) have been used to refine candidate CNVs by modeling read-depth signals as images, as seen in tools like CNV-espresso [16], imageCNV (Min et al., 2021), and DeepCNV [18]. These CNN-based methods typically rely on a relatively small subset of experimentally validated CNVs as gold standard training data, from which genomic features are learned to build supervised models for CNV identification in the new samples.

However, important challenges remain despite deep learning-based CNV detection strategies. First, CNN-based methods rely heavily on image recognition and are therefore prone to domain shift, with performance degrading when they are applied to datasets that differ substantially from the training data [19]. In addition, transforming one-dimensional read-depth signals into images and subsequently back into numerical representations can lead to loss of information, particularly for CNVs with weak signals. As a result, CNN-based approaches may struggle to generate reliable CNV calls when the dataset is not similar in structure or scale from the training dataset. Second, accurate supervised learning requires a large, validated variant set to more accurately capture the features of true CNVs. Existing DL methods use various strategies to construct “ground truth” CNVs, including human expert labeling with manual visualization [18], consensus calls from multiple tools [15], or CNV discoveries from matched whole-genome sequencing (WGS) experiments as ground truth [20]. However, these approaches are either subjective or difficult to scale to large datasets, and the results may be unreliable with the low-quality ground truth derived from other sequencing platforms.

To enhance model generalizability and the validity of CNV detection, we developed a novel deep learning framework, CN-RNN (<u>C</u>opy <u>N</u>umber variants detection with <u>R</u>ecurrent <u>N</u>eural <u>N</u>etworks). CN-RNN leverages Recurrent Neural Networks (RNNs) [21] to model read depth as an ordered genomic sequence, preserving local and long-range genomic dependencies. In addition, we utilized a publicly available parent-child trio dataset to construct a high-confidence gold standard CNV set based on the Mendelian rule of inheritance. By leveraging the dependent data structure across the genome and more reliable training labels, CN-RNN significantly enhances the accuracy of CNV detection. To rigorously assess accuracy and generalizability, we benchmarked CN-RNN against traditional and deep learning CNV detection methods on three independent datasets that differed in disease context, sequencing platform, and the construction of ground-truth CNVs (Mendelian inheritance from trio WES, experimental validation, and matched whole-genome sequencing), providing a more stringent evaluation than single-dataset comparisons. CN-RNN provides an accurate and computationally efficient approach for CNV detection in large-scale WES studies and supports broader clinical application of CNV analysis.

## Results

### Overview of CN-RNN

The schematic workflow of CN-RNN is shown in Figure 1, with details described in the <u>Methods</u> section. Briefly, CN-RNN was built as a deep learning-based tool that combined an existing high-sensitivity CNV caller with a deep learning model to capture genomic features to remove potential false positives. During training, CNV profiling was conducted by CORRseq [10] as a highly sensitive CNV caller. Ground truth CNVs were constructed using parent-offspring trio data from the Autism Sequencing Consortium (ASC) [22] to generate high-confidence positive and negative training labels. Using a hybrid neural network architecture, CN-RNN integrated sequential exon-level read depth with region-level genomic information to learn features that distinguished true CNVs from false positives via a supervised learning approach. The neural network architecture consisted of: (1) a bidirectional long short-term memory (BiLSTM) branch that encoded and learned the dependent sequential read-depth signal, and (2) a multi-layer perceptron (MLP) branch that encoded the genomic metadata embedded in each potential CNV region that was defined by the preliminary CNV caller (i.e., CORRseq). The representations learned from these branches were combined to generate high-confidence CNV predictions in the testing data.

**Figure 1.**
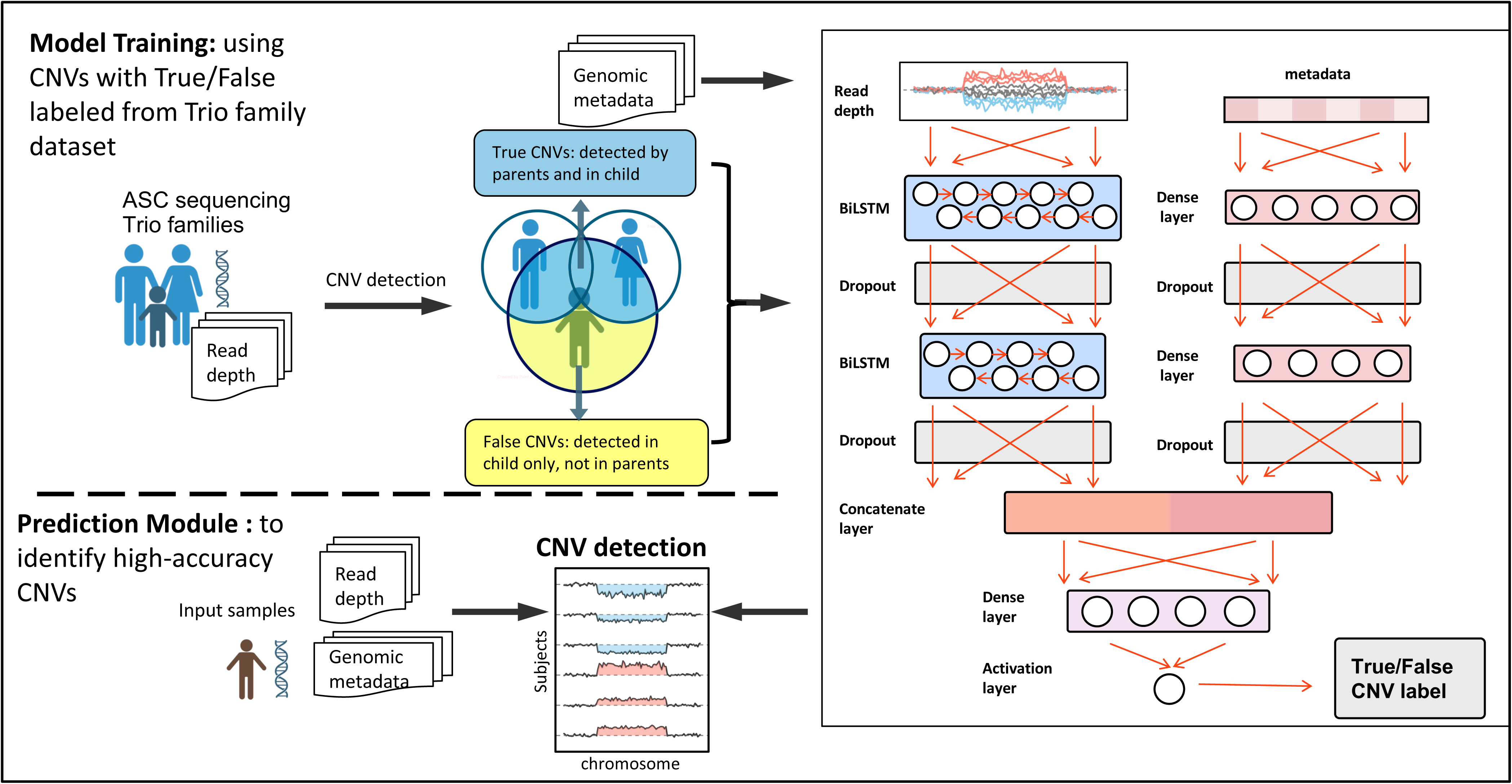
Overview of CN-RNN framework. During model training with Autism Sequencing Consortium (ASC) trio families, candidate CNVs were first detected from whole-exome sequencing (WES) data. CNVs detected in both the child and at least one parent were labeled as true CNVs, whereas CNVs detected only in the child were labeled as false CNVs based on the Mendelian rule of inheritance. Exon-level read-depth signals and region-level genomic metadata were used as model inputs. Read depth sequences were processed through a stacked bidirectional long short-term memory (BiLSTM) branch, while genomic metadata were encoded using a multi-layer perceptron (MLP) branch. Features from both branches were concatenated and passed through dense and activation layers to generate a binary true/false CNV prediction. In the prediction module, the trained CN-RNN model was applied to new samples to generate high-confidence CNV calls.

### CN-RNN achieves high accuracy in WES-based germline CNV detection with the ASC dataset

To rigorously develop and evaluate CN-RNN, we used the dataset from the Autism Sequencing Consortium (ASC) [22, 23] for model training and testing. Using the Mendelian rule of inheritance (<u>Methods</u>), high-confidence CNV labels were constructed using the identified CNVs from the offspring samples as ground truth. After pre- and post-CNV-calling quality control steps for all the samples (n = 971 offspring) (<u>Methods</u>, Supplementary Figure 1), 14,729 deletions and 24,665 duplications were labeled as true CNVs, and 63,142 deletions and 28,091 duplications were labeled as false CNVs. From the training samples (n = 874, 90% of the offspring samples), the labeled CNVs were down-sampled to form a balanced training set composed of 50% true CNVs and 50% non-CNV segments (i.e., 40% false CNVs and 10% negative controls) (<u>Methods</u>, Supplementary Table 1). After the training process with these high-quality ground truth CNVs, we evaluated the CN-RNN framework on the held-out ASC testing dataset (n = 97, 10% of the offspring samples). We compared CN-RNN against DL methods including CNV-espresso [16], DECoNT [24], and ECOLE [20], and widely used CNV callers, including CORRseq [10], EXCAVATOR2 [11], XHMM [12], CoNIFER [13], and CODEX2 [14].

As shown in Figure 2 and Supplementary Table 4, CN-RNN achieved the highest F1 score among all methods for both deletion and duplication detection. For deletions, CN-RNN reached an F1 score of 71%, representing a 2.2-fold improvement over DECoNT (32%, second-best method). For duplications, CN-RNN reached an F1 score of 73%, representing a 1.7-fold improvement over CNV-espresso (42%, second-best method). Deep learning methods (CN-RNN, CNV-espresso, DECoNT, ECOLE) overall outperformed traditional statistical tools (EXCAVATOR2, XHMM, CoNIFER, CODEX2) for both deletions and duplications. Benchmarked against CORRseq, CN-RNN’s precision increased from 27% to 72%, resulting in an F1 score increasing from 42% to 71% for deletions. Similarly, CN-RNN’s precision increased from 31% to 72%, resulting in an F1 score increasing from 47% to 73% for duplications. Analysis across CNV lengths revealed that the performance gain was most pronounced in short CNVs (5-10 exons). Such improvement for short CNVs was important because short CNVs produced weaker signals and had historically been hard for most WES CNV callers to detect. Notably, the dual-branch CN-RNN architecture processed genomic metadata in parallel with read-depth signals, providing auxiliary information that helped the identification of short CNVs. In summary, CN-RNN outperformed all competing methods in overall accuracy and exhibited the most pronounced performance gain in short CNVs.

**Figure 2.**
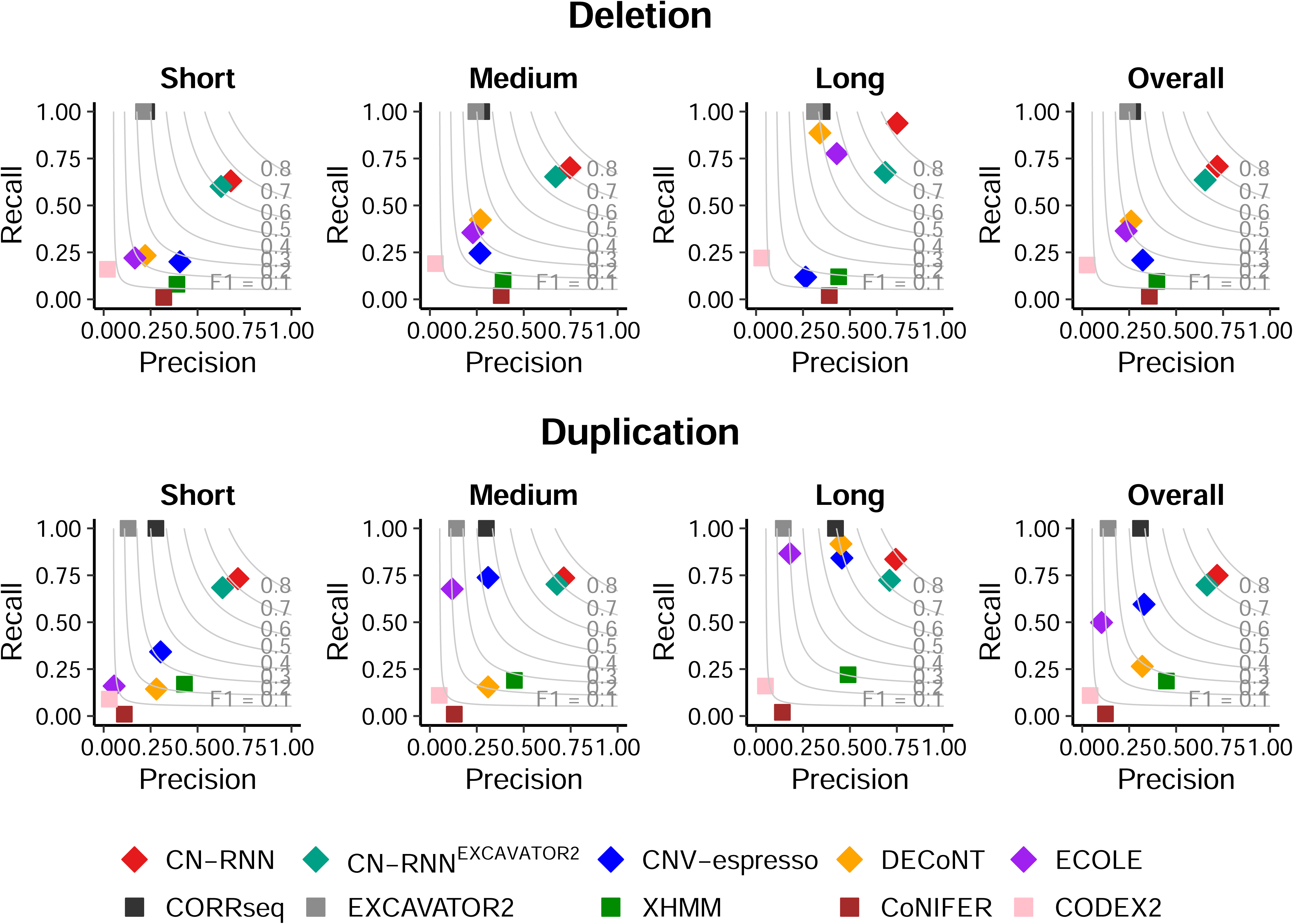
Performance on the Autism Sequencing Consortium (ASC) testing dataset. Precision-recall scatter plots are shown for CN-RNN, CN-RNN^EXCAVATOR2^ (CN-RNN variant model trained on EXCAVATOR2-derived training set), and existing CNV detection methods including CNV-espresso, DECoNT, ECOLE, CORRseq, EXCAVATOR2, XHMM, CoNIFER, and CODEX2 on the Autism Sequencing Consortium (ASC) trio testing set. Plots are shown for deletions and duplications, stratified by CNV length: short (5-10 exons), medium (11-40 exons), long (>40 exons) and overall (all CNV lengths pooled). Gray contour lines indicate constant F1 scores.

To further evaluate the flexibility and robustness of CN-RNN, we also retrained the same hybrid neural network architecture after an alternative CNV caller, EXCAVATOR2 [11], instead of CORRseq for copy number estimation in the training process. Using the new training CNV labels, we obtained a variant model, denoted as CN-RNN^EXCAVATOR2^. Compared to the original CN-RNN training set (i.e., using CORRseq as the preliminary CNV caller), the CN-RNN^EXCAVATOR2^ training set was smaller (Supplementary Table 2). Direct comparison between the original CN-RNN and CN-RNN^EXCAVATOR2^ showed better performance of the original CN-RNN in the ASC testing dataset (Figure 2 and Supplementary Table 4). This difference likely reflects the higher recall of CORRseq, especially in detecting short CNVs, which produced a larger and more diverse training set with richer signal patterns for the model to learn from. Nevertheless, CN-RNN^EXCAVATOR2^ successfully improved over EXCAVATOR2 by increasing the F1 score from 39% to 65% for deletions and from 24% to 68% for duplications, demonstrating the feasibility of implementing the CN-RNN hybrid neural network framework with alternative preliminary CNV callers.

In addition, CN-RNN demonstrated favorable computational efficiency compared to other methods. Table 1 summarizes the average computation time per ASC sample for all methods. The DL methods (CN-RNN, CNV-espresso, DECoNT, ECOLE) overall had longer computational time than non-DL methods. Among the DL methods, CN-RNN achieved the highest computational efficiency despite its dual-branch and multi-layer design, running faster than CNV-espresso (convolutional neural network, CNN), DECoNT (single-layer RNN), or ECOLE (transformer-based). This efficiency was achieved through optimized preprocessing and parallel computing (Supplementary Notes).

**Table 1.** Computational time comparison. Average computational time per sample for CN-RNN and competing CNV detection methods, including preprocessing, CNV calling, and model inference, evaluated under the same computational environment. The sample was from the ASC dataset. CN-RNN^EXCAVATOR2^ denotes the variant model of CN-RNN trained with EXCAVATOR2 as preliminary CNV caller.

| Method | Computational time per sample (seconds) |
| --- | --- |
| CN-RNN | 35 |
| CN-RNN<sup>EXCAVATOR2</sup> | 35 |
| CNV-espresso | 45 |
| DECoNT | 49 |
| ECOLE | 2,400 |
| CORRseq | 30 |
| EXCAVATOR2 | 24 |
| XHMM | 51 |
| CoNIFER | 205 |
| CODEX2 | 130 |

### External validation on the Pediatric Cardiac Genomics Consortium (PCGC) dataset

To assess generalizability beyond the training cohort and sequencing platforms, CN-RNN was evaluated on an independent dataset from the Pediatric Cardiac Genomics Consortium (PCGC) [25], where candidate CNVs were experimentally validated. For this dataset, we evaluated detection performance on a subset of 18 validated deletions and 4 duplications from 21 samples. Supplementary Figure 2 shows the preprocessing and quality control details. Although the number of validated CNVs was relatively small, external validation using experimentally confirmed CNVs provided a more rigorous assessment compared to the *in silico* labeling approaches.

As the PCGC dataset provided only a set of experimentally validated CNV events, non-CNV regions were not available; performance was therefore assessed primarily by recall rate (<u>Methods</u>). We compared CN-RNN against CNV-espresso, DECoNT, ECOLE, EXCAVATOR2, XHMM, CoNIFER, and CODEX2. As shown in Figure 3, CN-RNN achieved the highest recall for both deletions and duplications, successfully detecting 14 out of 18 deletions (recall = 78%) and 3 of 4 duplications (recall = 75%). Detailed CNV validation results by CN-RNN are provided in Supplementary Table 5. For deletions, the second-best method, XHMM recovered 12 of 18 deletions (recall = 67%). For duplications, ECOLE recovered 3 of 4 duplications (recall = 75%), while most other methods detected only 2 of 4 duplications (recall = 50%). CN-RNN performance was consistent across recurrent and rare CNV categories, it identified 13 of 16 rare CNVs (recall = 81%) and 4 of 6 recurrent ones (recall = 67%).

**Figure 3.**
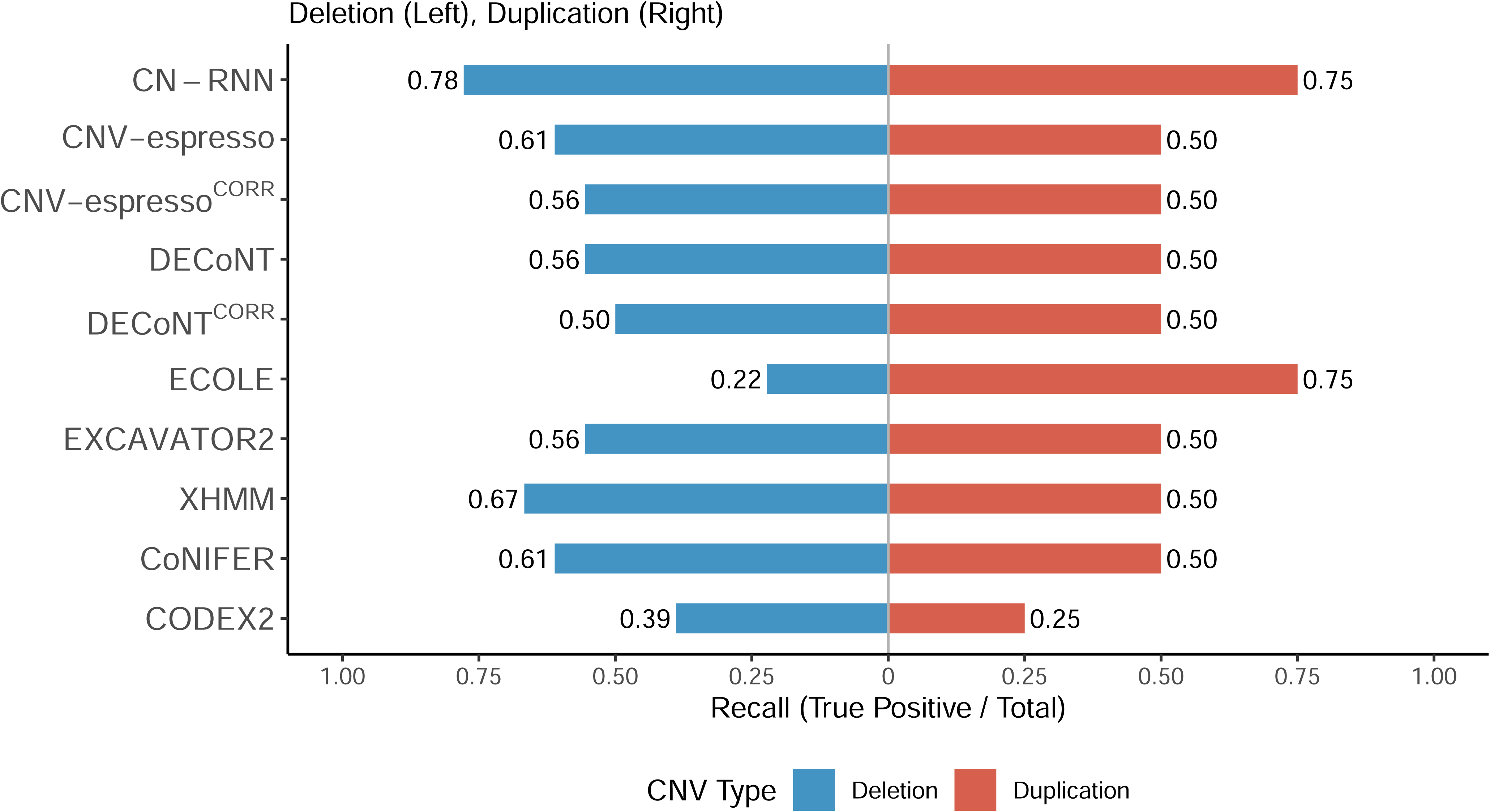
Performance on experimentally validated CNVs in the Pediatric Cardiac Genomics Consortium (PCGC) dataset. Recall of CNV detection methods on the PCGC dataset is shown as a bar chart for 18 deletions (left) and 4 duplications (right). CNV-espresso and DECoNT used XHMM as the initial CNV caller, whereas CNV-espresso^CORR^ and DECoNT^CORR^ used CORRseq as the initial CNV caller.

Because CNV-espresso and DECoNT functioned as post-calling confirmation methods and relied on XHMM as their primary CNV callers, the evaluation did not allow a parallel comparison with our method on the neural network architecture alone, because the two tools differ from CN-RNN in both CNV preliminary callers and deep learning architectures. We therefore equipped these two tools with CORRseq as the preliminary CNV caller to benchmark against CN-RNN on their deep learning components independently. These two frameworks were therefore denoted as CNV-espresso^CORR^ and DECoNT^CORR^, which showed lower recall than their original XHMM-based versions (Figure 3). CN-RNN outperformed both, further demonstrating the advantage of our DL framework.

Importantly, the performance of CN-RNN was achieved without any cohort-specific calibration or parameter tuning, indicating that the signal patterns learned from the ASC trio dataset were transferable to independent sequencing platforms. The PCGC dataset differed substantially from the ASC training cohort in sequencing platforms, exon capture kits, and cohort composition. Despite these cross-cohort differences, CN-RNN successfully identified most experimentally validated CNVs, indicating robustness to platform-dependent bias and batch effects.

### External validation on the 1000 Genomes Project dataset

Finally, we evaluated model performance on samples from the 1000 Genomes Project [26], where matched whole-genome sequencing (WGS)-derived CNV calls served as a semi-ground-truth reference set for cross-platform concordance assessment. Supplementary Figure 3 shows the preprocessing and quality control details. As shown in Figure 4 and Supplementary Table 6, the overall CNV detection performance was low across all evaluated methods for both deletions and duplications, except for CN-RNN and ECOLE. For deletions, CN-RNN achieved the highest F1 score (24%), doubling the performance of the second-best method, CNV-espresso (12%). For duplications, CN-RNN and ECOLE both achieved an F1 score of 20%. CN-RNN’s relatively higher F1 score was driven primarily by its higher recall compared to other methods. We also observed that CN-RNN maintained a high recall (78% and 83% for deletions and duplications) comparable to rates achieved in the ASC and PCGC evaluations, especially in medium and long CNVs (Supplementary Table 6 and Supplementary Figure 4). However, precision (14% and 11% for deletions and duplications, respectively) was lower than that observed in the ASC testing dataset.

**Figure 4.**
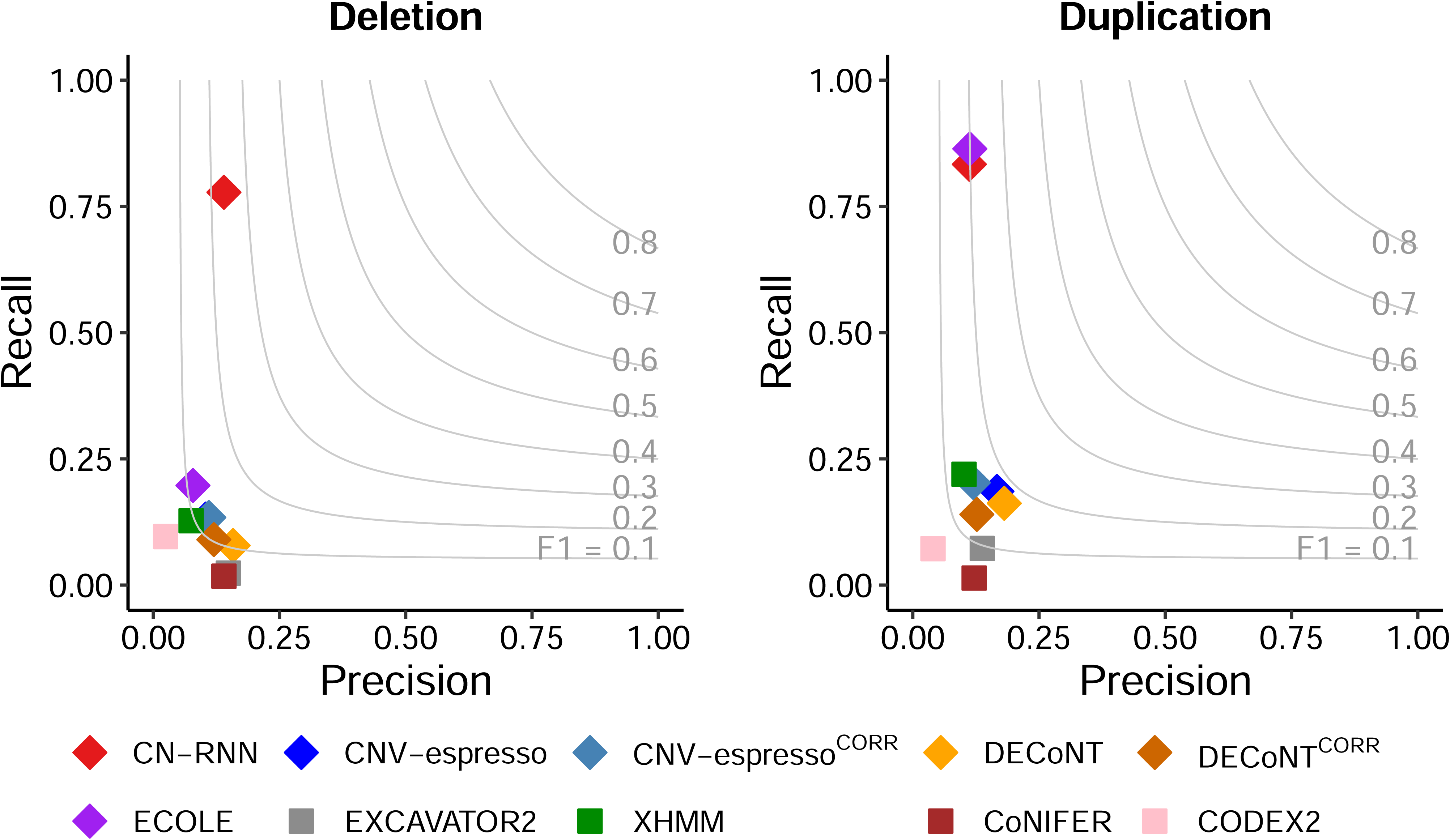
Overall Performance on the 1000 Genomes Project dataset. Precision-recall scatter plots are shown for CN-RNN and competing methods on the 1000 Genomes Project WES dataset, summarized across all CNV lengths. Results are shown separately for deletions (left) and duplications (right). CNV-espresso and DECoNT used XHMM as the initial CNV caller, whereas CNV-espresso^CORR^ and DECoNT^CORR^ used CORRseq as the initial CNV caller.

The reduced overall performance was expected, as the WGS-derived CNVs in the 1000 Genomes Project dataset differed from the CN-RNN training set in both CNV size distribution and breakpoint resolution. Specifically, the median CNV lengths detected by CNVnator from WGS data were 13.1 kb for deletions and 12.6 kb for duplications, whereas the corresponding values detected by CN-RNN from WES data were 188 kb and 286 kb. In addition, the WGS-derived CNV set had base-pair-level resolution, whereas WES-based CNV callers (except ECOLE) used exons as the fundamental detection units and therefore operated at a lower resolution in CNV detection. CN-RNN was trained on GRCh37-aligned WES data but evaluated on 1000 Genomes samples which were GRCh38-aligned; this cross-build application may have introduced differences in exon coordinates and read-depth profiles that further contributed to the reduced precision. Despite these challenges, the higher recall achieved by CN-RNN resulted in comparatively strong overall performance among the evaluated methods.

## Discussion

In this study, we developed CN-RNN, a deep learning method for CNV detection from WES read-depth signals. The contribution of this work is primarily reflected in three aspects. First, we constructed training labels from the Mendelian rule of inheritance in a trio-structured WES cohort, a strategy that circumvented the need for experimental validation or human expert curation and enabled large-scale label generation for deep learning. Second, we designed a novel dual-branch neural network architecture that combined a read-depth processing branch using BiLSTM with a parallel genomic metadata branch. This design mitigated the systematic capture biases (e.g., exon size, GC content, mappability) that limit purely depth-based callers. Third, we benchmarked CN-RNN against existing callers on three independent datasets, each with different ground-truth construction criteria. Across all three evaluation settings, CN-RNN achieved the highest CNV detection accuracy among the methods compared, with the largest gains observed for short CNVs while remaining competitive in computational cost.

A persistent obstacle to applying deep learning methods to CNV detection has been the lack of large-scale CNV sets with validated training labels. Experimentally validated CNV sets and expert-curated gold-standard call sets are typically small and limited to specific disease contexts. We made use of the trio design of the ASC cohort to derive training labels from the Mendelian rule of inheritance, building on principles established in family-based CNV studies [22]. By applying this strategy to approximately 3,100 trio WES samples, we generated a novel labeled training set substantially larger than the curated sets used by prior WES CNV deep learning methods, without requiring additional sequencing or molecular assays.

Furthermore, CN-RNN used a dual-branch architecture, which processed the read-depth signal in a BiLSTM branch and genomic metadata in a parallel MLP branch before their representations were combined. The BiLSTM branch modeled exon-level read depth as an ordered genomic sequence, naturally capturing contextual dependencies across neighboring exons without introducing artificial spatial structures. The metadata branch acted as an auxiliary information channel that mitigated read-depth biases arising from the sequencing process, such as those from GC content and mappability. This dual-branch design yielded the largest gains for short CNVs (5-10 exons) in our benchmarks, where the read-depth signal alone was weak and hence the benefit from genomic-context information was amplified. More broadly, this result suggests that integrating genomic context alongside primary sequencing signals may be a useful strategy for other CNV detection tasks.

CN-RNN was evaluated on three datasets with different ground-truth sources: ASC (inheritance-derived labels from trio WES), PCGC (experimentally validated CNVs in an independent disease cohort), and 1000 Genomes Project (CNVnator WGS calls on matched samples). Across all three settings, CN-RNN consistently performed well, demonstrating the practical adequacy of the framework. Importantly, the high recall rate in PCGC and F1 score gain in 1000 Genomes Project were achieved without retraining or parameter tuning on the external datasets, indicating that CN-RNN learned transferable signal patterns of WES-derived CNVs rather than overfitting to a specific dataset or cohort.

Still, several limitations should be acknowledged in this study. First, each of the three validation datasets has limitations in defining its ground truth. The ASC ground truth depended on the Mendelian rule of inheritance and could not distinguish between *de novo* CNV mutations and false positive calls. The PCGC validation set comprised only 22 experimentally validated CNVs aggregated from previously published PCGC studies, which was a small set and subject to selection bias toward CNVs of clinical interest. The 1000 Genomes Project evaluation used CNVnator WGS calls as a semi-ground-truth rather than an absolute reference. Second, CN-RNN relied on CORRseq for initial breakpoint discovery and candidate region generation. Consequently, the recall of the overall framework was partially constrained by the discovery capability of CORRseq. We also evaluated EXCAVATOR2 as an alternative preliminary caller, but CORRseq was preferred because of its higher sensitivity. Nevertheless, our deep learning architecture is easily adapted to other callers in future studies. Third, extremely short CNVs spanning fewer than five exons fell outside the current scope. Extending CN-RNN to this range will be a natural direction for future work and would likely require denser per-exon feature representations in the training process. Another direction is to develop a CN-RNN variant that performs CNV detection independently of CORRseq or other preliminary callers, which would remove the recall ceiling imposed by the initial CNV calling step. In addition, although CN-RNN was motivated by the challenges in germline CNV detection, the approach can be readily extended to study somatic copy number changes in disease contexts such as cancer.

## Methods

### Datasets

#### The Autism Sequencing Consortium (ASC)

To facilitate feature learning in the CN-RNN framework, we used 978 parent-child trio families with at least one child diagnosed with autism spectrum disorder and 10 control families (a total of 3,100 samples) from the Autism Sequencing Consortium (ASC) [22]. The ASC study aggregated WES data from more than 12,000 trio-structured samples (over 3,000 families) to investigate genetic risk factors for autism spectrum disorder. The data were sequenced on the Illumina HiSeq X Ten platform and downloaded from dbGaP (accession: phs000298.v4.p3). All sequenced reads were aligned to the GRCh37 human reference genome. The trio structure of the ASC dataset enabled systematic construction of high-confidence CNV labels based on the Mendelian rule of inheritance. We used 10 healthy families without disease history to serve as reference samples for read depth normalization.

#### The Pediatric Cardiac Genomics Consortium (PCGC)

To evaluate the generalizability of CN-RNN to experimentally validated variants, we benchmarked CN-RNN and other methods on an independent WES dataset from the Pediatric Cardiac Genomics Consortium (PCGC) study [25]. The PCGC is a multi-center prospective cohort designed to investigate the genetic basis of congenital heart defects (CHD) (dbGaP accession: phs000571.v7.p3). We selected a subset of WES samples sequenced on the Illumina HiSeq 2000 platform and aligned to the GRCh37 reference genome. This subset comprised 22 experimentally validated CNVs falling into two categories: six recurrent CNVs, defined as those occurring at known pathogenic loci in multiple unrelated individuals, including loci at 22q11.2 and 7q11.23, validated using multiplex ligation-dependent probe amplification (MLPA); and 16 rare CNVs, defined as those with a frequency < 0.01 in the cohort [16, 27, 28], validated by droplet digital PCR (ddPCR). In addition, 12 samples without validated CNVs were included as reference samples for read depth normalization.

#### The 1000 Genomes Project

The 1000 Genomes Project is a widely used benchmark dataset that provides matched WES and WGS data [26]. We analyzed 901 samples with WES data generated on Illumina Genome Analyzer II and Illumina HiSeq 2000 platforms and the NimbleGen SeqCap v3 capture kit. Matched WGS data for these samples were sequenced on the Illumina NovaSeq platform. High-confidence CNVs were identified from WGS data using CNVnator [29]. These calls (1,219,094 deletions, 654,659 duplications) were considered as semi-ground-truth for benchmarking the methods. Both WES and WGS reads were aligned to the GRCh38 reference genome.

### CN-RNN Method Details

#### Data processing and CNV profiling with the ASC dataset

With the 978 trio samples, we first calculated exon-level read counts from the aligned BAM files using Rsubread [30]. The resulting read count matrix was then preprocessed through extensive quality control and normalization steps including a median-based normalization approach from EXCAVATOR [31], scaling to pre-specified normal control samples and a smoothing procedure (Supplementary Notes, Supplementary Figure 1) [32]. After CNV calling with CORRseq [10], post-CNV calling quality control steps included (a) retaining those spanning between 5 and 120 exons and shorter than 5,000 kb, and (b) excluding samples with an excessive number of CNV calls (> 500). The significance level was set at 0.05 for CORRseq.

#### Construction of CNV training profile

To construct a high-confidence CNV call set for training, we applied the Mendelian rule of inheritance to identify true CNVs within the training data, a strategy previously applied in CNV-espresso for label assignment [16, 33]. For CNVs identified in offspring, those also detected in at least one parent were classified as true CNVs, whereas those detected only in the offspring but not in either parent were considered Mendelian errors and excluded from the true CNV set. Mendelian errors can arise from three possible mechanisms: (1) a *de novo* CNV in the offspring, (2) a false positive call in the offspring, or (3) a false negative call in a parent. *De novo* CNVs occur at a frequency of approximately 0.01-0.02 in the general population and 0.05-0.10 in individuals with autism [34].

In the training dataset (n = 874, 90% of samples from the ASC dataset), we constructed a balanced CNV training set with a combination of (1) true CNVs (50%), (2) false CNVs from Mendelian errors (40%), and (3) negative controls representing normal diploid regions (10%) (Supplementary Table 1). The negative controls were randomly sampled from genomic locations where no CNVs were observed. To minimize selection bias, the chromosomal locations and segment sizes of these negative controls were matched to those of the true CNVs. False CNVs and negative control segments were grouped as non-CNV segments for model training, resulting in a binary classification task distinguishing true CNVs from non-CNV regions. During down-sampling, we ensured that comparable numbers of training examples were distributed across various CNV lengths to allow the model to learn across different length ranges.

#### Problem formulation

For the CN-RNN deep learning model, we assumed there were *n_del_*, *n_dup_* deletion and duplication events within the training CNV set. Let *n_all_* = *n_del_* + *n_dup_* denote the total number of CNV events. For classification, the main inputs for each event *X*^(*i*)^, *i* ∈ {1,2,…, *n_all_*} were represented by a feature set *F*^(*i*)^ = {*R*^(*i*)^,*M*^(*i*)^}. *R*^(*i*)^ was a sequential read depth vector capturing the normalized exon-level signal (*log2R*, i.e., the log-2 ratio of normalized read depth) spanning the *i*-th CNV, preserving local genomic context. *M*^(*i*)^ was a genomic metadata feature vector summarizing region-level characteristics, including median GC content, median mappability, number of exons per CNV region, CNV length in kilobases (log10-transformed), mean *log2R*, standard deviation of *log2R*, and chromosome ID. These features were used as input to the MLP branch of the CN-RNN model. A detailed description of these features is provided in Supplementary Table 3. Data normalization and smoothing procedures are detailed in Supplementary Notes.

For each event, we assigned a binary label 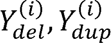 ∈ {0,1}, where *Y*^(*i*)^ =1 denoted a true CNV and *Y*^(*i*)^ =0 denoted a non-CNV. Because deletions and duplications displayed distinct read depth patterns, we trained two independent classifiers: 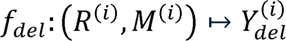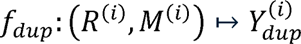. The two models for deletions and duplication did not share parameters and were optimized separately on each subset, but they shared the same underlying neural network architecture.

#### CN-RNN model architecture

We employed a hybrid neural network consisting of two parallel branches to integrate sequential read-depth signals and region-level genomic metadata (Figure 1). The exon-level read depth sequence *R*^(*i*)^ (for the *i*-th CNV event) was fed into a two-layer bidirectional long short-term memory (BiLSTM) network. The two stacked BiLSTM layers contained 64 and 32 hidden units, respectively. To mitigate overfitting, a dropout layer (rate = 0.5) was applied after each BiLSTM layer, and an L2 kernel regularization term (*λ* = 0.01) was incorporated in the second BiLSTM layer. This branch was designed to capture local decreases or increases in read depth that characterize deletions or duplications, while also modeling the contextual dependencies provided by neighboring exons across the CNV segments. In parallel, the metadata branch encoded the genomic metadata vector *M*^(*i*)^. This branch consisted of two dense layers with 64 and 32 neurons, each followed by ReLU activation and dropout (rate = 0.5). Outputs from the sequential and metadata branches were concatenated and passed through two additional dense layers (64 and 32 neurons, ReLU activation, dropout = 0.5), followed by a final sigmoid layer to classify each candidate event as a true CNV or non-CNV.

The formulation of CN-RNN can be summarized as follows:

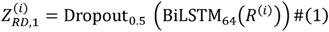

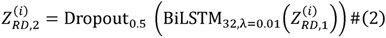

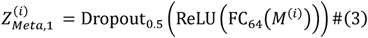

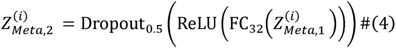

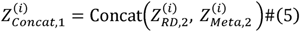

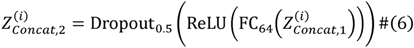

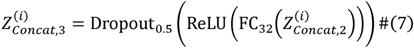

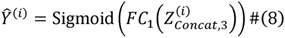

Here, FC*_k_* denotes a fully connected layer with *k* neurons, BiLSTM*_k_* denotes a bidirectional LSTM with *k* hidden units, Dropout*_r_* denotes dropout with rate *r*, Concat(·,·) denotes concatenation of feature vectors, and Sigmoid(·) is the sigmoid activation function, which maps the final output to a probability for binary classification.

#### CN-RNN model training and computational details

CN-RNN was optimized using the Adam optimizer (learning rate = 0.0001) with a binary cross-entropy loss function. Five-fold cross-validation was performed on the training set to ensure model stability and prevent overfitting. Each model was trained for 40 epochs with a batch size of 120 (chosen to balance computational efficiency and gradient stability), with early stopping applied if the validation loss did not decrease for five consecutive epochs. CN-RNN training and evaluation, along with benchmarking of all comparison methods, were performed on the HiPerGator high-performance computing cluster at the University of Florida, with identical computational resources. Each job was allocated 16 CPU cores and 120 GB of RAM on nodes powered by AMD EPYC 9655P processors. The 120 GB allocation was set as a uniform upper bound to accommodate the most memory-intensive method among those benchmarked. CN-RNN’s observed peak memory usage was approximately 17 GB. Among the benchmarked methods, ECOLE’s runtime was substantially longer due to its transformer-based architecture and base-pair-level resolution, which could potentially be accelerated by GPU rather than CPU execution. To ensure a fair comparison, all methods were run on CPUs under a unified computational environment.

#### Performance evaluation with three datasets

##### Evaluation on the ASC trio dataset

Performance was assessed on the 10% (n = 97) held-out testing set using inheritance-filtered ground-truth labels derived from CORRseq and trio-based Mendelian criteria. A predicted CNV in the offspring was considered a true CNV if it overlapped with a CNV in the inheritance-derived ground-truth set by more than 20% of its genomic length.

##### Evaluation on the PCGC dataset

The PCGC dataset was used to test CN-RNN’s ability to identify experimentally validated CNVs (n = 22) across different sequencing platforms and exon capture designs. A prediction was counted as recovery of a validated CNV when it shared the same CNV type and overlapped the validated event by more than 20% of its genomic length. Because genome-wide negative labels were unavailable, performance was summarized by recall.

##### Evaluation on the 1000 Genomes Project dataset

Performance on the 1000 Genomes Project dataset (n = 901) was evaluated by comparing WES-based CNV predictions against WGS-derived CNVs identified by CNVnator, which were treated as a semi-ground truth reference. A predicted CNV was considered a match when it overlapped a WGS CNV of the same type by more than 1% of its genomic length, a relaxed threshold chosen to accommodate breakpoint resolution differences between WES and WGS. Because WGS-derived CNVs represented a proxy rather than experimentally validated ground truth, these metrics reflected relative WES-WGS concordance and were used primarily for benchmarking CN-RNN against existing methods, rather than for absolute accuracy estimation.

Across all three datasets, CN-RNN was benchmarked against CNV-espresso, DECoNT, ECOLE, CORRseq, EXCAVATOR2, XHMM, CoNIFER, and CODEX2. All analyses were restricted to autosomal chromosomes to ensure consistency across CNV callers. For methods providing numerical copy number estimates (e.g., EXCAVATOR2, CODEX2), CNV status was classified as a deletion if the estimated copy number was < 2 and as a duplication if it was > 2. Performance was quantified by precision, recall, and F1 score, computed separately for duplications and deletions and stratified by CNV length to assess robustness across CNV sizes.

Precision, recall, and F1 score were defined as follows:

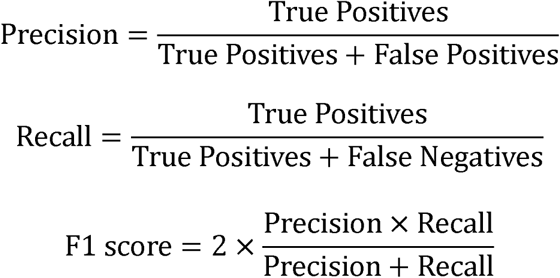

## Data Availability

The datasets used to train and evaluate CN-RNN are subject to their respective data access policies. Whole-exome sequencing data from the Autism Sequencing Consortium (ASC) are available from the Database of Genotypes and Phenotypes (dbGaP) under accession phs000298.v4.p3 upon approved data access request [22, 23]. Whole-exome sequencing data from the Pediatric Cardiac Genomics Consortium (PCGC) are generated under the auspices of the National Heart, Lung, and Blood Institute’s Bench to Bassinet Program (https://benchtobassinet.com). The data are available from dbGaP under accession phs000571.v7.p3 upon approved data access request [25]. The 1000 Genomes Project whole-exome and whole-genome sequencing data used for validation are publicly available from the International Genome Sample Resource (IGSR) at https://ftp.1000genomes.ebi.ac.uk/vol1/ftp/data_collections/1000_genomes_project/data/ [26]. Samples from HG00096 to HG02356 were selected for our study.

CN-RNN is implemented and released on GitHub (https://github.com/FeifeiXiao-lab/CN-RNN). The pipeline accepts aligned WES BAM files as input. The repository includes the trained model, pre-processing scripts, and all Python and R scripts used to apply CN-RNN to new WES samples.

## Key Points

- CN-RNN is a novel deep-learning-based CNV caller for WES data that introduces a dual-branch neural network combining a bidirectional long short-term memory (BiLSTM) branch for exon-level read-depth sequences with a parallel multi-layer perceptron (MLP) branch for region-level genomic metadata, mitigating systematic capture biases that limit purely depth-based methods and yielding the largest accuracy gains for short CNVs.
- CN-RNN constructs high-confidence CNV training labels at large scale by applying the Mendelian rule of inheritance to parent-child trio WES data, removing the dependence on small experimentally validated CNV sets or expert-curated gold-standard calls that has constrained prior deep-learning callers.
- Benchmarked across three independent datasets spanning distinct disease contexts, sequencing platforms, and ground-truth construction strategies (trio inheritance, experimental validation, and matched WGS), CN-RNN achieved the highest F1 and recall among competing methods while remaining computationally efficient and transferable without cohort-specific retraining.

## Reference

[1] Zarrei M, MacDonald JR, Merico D, Scherer SW. A copy number variation map of the human genome. Nature Reviews. Genetics. 2015;16(3):172–183. doi:10.1038/nrg3871

[2] Horpaopan S, Spier I, Zink AM, Altmüller J, Holzapfel S, Laner A, Vogt S, Uhlhaas S, Heilmann S, Stienen D, et al. Genome-wide CNV analysis in 221 unrelated patients and targeted high-throughput sequencing reveal novel causative candidate genes for colorectal adenomatous polyposis. International Journal of Cancer. 2015;136(6):E578–E589. doi:10.1002/ijc.29215

[3] Kumaran M, Cass CE, Graham K, Mackey JR, Hubaux R, Lam W, Yasui Y, Damaraju S. Germline copy number variations are associated with breast cancer risk and prognosis. Scientific Reports. 2017;7(1):14621. doi:10.1038/s41598-017-14799-7

[4] Lander J, Ware SM. Copy Number Variation in Congenital Heart Defects. Current Genetic Medicine Reports. 2014;2(3):168–178. doi:10.1007/s40142-014-0049-3

[5] Alhazmi S, Alzahrani M, Farsi R, Alharbi M, Algothmi K, Alburae N, Ganash M, Azhari S, Basingab F, Almuhammadi A, et al. Multiple Recurrent Copy Number Variations (CNVs) in Chromosome 22 Including 22q11.2 Associated with Autism Spectrum Disorder. Pharmacogenomics and Personalized Medicine. 2022;15:705–720. doi:10.2147/PGPM.S366826

[6] Castellani CA, Awamleh Z, Melka MG, O’Reilly RL, Singh SM. Copy Number Variation Distribution in Six Monozygotic Twin Pairs Discordant for Schizophrenia. Twin Research and Human Genetics. 2014;17(2):108–120. doi:10.1017/thg.2014.6

[7] Gabrielaite M, Torp MH, Rasmussen MS, Andreu-Sánchez S, Vieira FG, Pedersen CB, Kinalis S, Madsen MB, Kodama M, Demircan GS, et al. A Comparison of Tools for Copy-Number Variation Detection in Germline Whole Exome and Whole Genome Sequencing Data. Cancers. 2021;13(24):6283. doi:10.3390/cancers13246283

[8] Pirooznia M, Goes FS, Zandi PP. Whole-genome CNV analysis: advances in computational approaches. Frontiers in Genetics. 2015 [accessed 2025 June 12];6. https://www.frontiersin.org/journals/genetics/articles/10.3389/fgene.2015.00138/full. doi:10.3389/fgene.2015.00138

[9] Tan R, Wang Y, Kleinstein SE, Liu Y, Zhu X, Guo H, Jiang Q, Allen AS, Zhu M. An Evaluation of Copy Number Variation Detection Tools from Whole-Exome Sequencing Data. Human Mutation. 2014;35(7):899–907. doi:10.1002/humu.22537

[10] Qin F, Luo X, Cai G, Xiao F. Shall genomic correlation structure be considered in copy number variants detection? Briefings in Bioinformatics. 2021;22(6):bbab215. doi:10.1093/bib/bbab215

[11] D’Aurizio R, Pippucci T, Tattini L, Giusti B, Pellegrini M, Magi A. Enhanced copy number variants detection from whole-exome sequencing data using EXCAVATOR2. Nucleic Acids Research. 2016;44(20):e154. doi:10.1093/nar/gkw695

[12] Fromer M, Purcell SM. Using XHMM Software to Detect Copy Number Variation in Whole-Exome Sequencing Data. Current Protocols in Human Genetics. 2014;81(1):7.23.1–7.23.21. doi:10.1002/0471142905.hg0723s81

[13] Krumm N, Sudmant PH, Ko A, O’Roak BJ, Malig M, Coe BP, Quinlan AR, Nickerson DA, Eichler EE. Copy number variation detection and genotyping from exome sequence data. Genome Research. 2012;22(8):1525–1532. doi:10.1101/gr.138115.112

[14] Jiang Y, Wang R, Urrutia E, Anastopoulos IN, Nathanson KL, Zhang NR. CODEX2: full-spectrum copy number variation detection by high-throughput DNA sequencing. Genome Biology. 2018;19(1):202. doi:10.1186/s13059-018-1578-y

[15] Pounraja VK, Jayakar G, Jensen M, Kelkar N, Girirajan S. A machine-learning approach for accurate detection of copy number variants from exome sequencing. Genome research. 2019;29(7):1134–1143. doi:10.1101/gr.245928.118

[16] Tan R, Shen Y. Accurate *in silico* confirmation of rare copy number variant calls from exome sequencing data using transfer learning. Nucleic Acids Research. 2022;50(21):e123–e123. doi:10.1093/nar/gkac788

[17] Min Q, Li X, Wang R, Ming H, Wang K, Hao X, Wang Y, Zhan Q. Accurate detection of CNV based on single-nucleotide variants recalibration and image classification from whole genome sequencing. Medicine in Omics. 2021;1:100002. doi:10.1016/j.meomic.2020.100002

[18] Glessner JT, Hou X, Zhong C, Zhang J, Khan M, Brand F, Krawitz P, Sleiman PMA, Hakonarson H, Wei Z. DeepCNV: a deep learning approach for authenticating copy number variations. Briefings in Bioinformatics. 2021;22(5):bbaa381. doi:10.1093/bib/bbaa381

[19] Takahashi S, Sakaguchi Y, Kouno N, Takasawa K, Ishizu K, Akagi Y, Aoyama R, Teraya N, Bolatkan A, Shinkai N, et al. Comparison of Vision Transformers and Convolutional Neural Networks in Medical Image Analysis: A Systematic Review. Journal of Medical Systems. 2024;48(1):84. doi:10.1007/s10916-024-02105-8

[20] Mandiracioglu B, Ozden F, Kaynar G, Yilmaz MA, Alkan C, Cicek AE. ECOLE: Learning to call copy number variants on whole exome sequencing data. Nature Communications. 2024;15(1):132. doi:10.1038/s41467-023-44116-y

[21] Schuster M, Paliwal KK. Bidirectional recurrent neural networks. IEEE Transactions on Signal Processing. 1997;45(11):2673–2681. doi:10.1109/78.650093

[22] Buxbaum JD, Daly MJ, Devlin B, Lehner T, Roeder K, State MW, Autism Sequencing Consortium. The autism sequencing consortium: large-scale, high-throughput sequencing in autism spectrum disorders. Neuron. 2012;76(6):1052–1056. doi:10.1016/j.neuron.2012.12.008

[23] Autism Sequencing Consortium. [dataset] Whole-exome sequencing data from the Autism Sequencing Consortium. 2012. https://www.ncbi.nlm.nih.gov/projects/gap/cgi-bin/study.cgi?study_id=phs000298.v4.p3

[24] Özden F, Alkan C, Çiçek AE. Polishing copy number variant calls on exome sequencing data via deep learning. Genome Research. 2022;32(6):1170–1182. doi:10.1101/gr.274845.120

[25] Pediatric Cardiac Genomics Consortium. [dataset] Whole-exome sequencing data from the Pediatric Cardiac Genomics Consortium. 2013. https://www.ncbi.nlm.nih.gov/projects/gap/cgi-bin/study.cgi?study_id=phs000571.v7.p3

[26] 1000 Genomes Project Consortium. [dataset] Whole-exome and whole-genome sequencing data from the 1000 Genomes Project. 2015. https://ftp.1000genomes.ebi.ac.uk/vol1/ftp/data_collections/1000_genomes_project/data/

[27] Glessner JT, Bick AG, Ito K, Homsy JG, Rodriguez-Murillo L, Fromer M, Mazaika E, Vardarajan B, Italia M, Leipzig J, et al. Increased Frequency of De Novo Copy Number Variants in Congenital Heart Disease by Integrative Analysis of Single Nucleotide Polymorphism Array and Exome Sequence Data. Circulation Research. 2014;115(10):884–896. doi:10.1161/CIRCRESAHA.115.304458

[28] Jin SC, Homsy J, Zaidi S, Lu Q, Morton S, DePalma SR, Zeng X, Qi H, Chang W, Sierant MC, et al. Contribution of rare inherited and de novo variants in 2,871 congenital heart disease probands. Nature genetics. 2017;49(11):1593–1601. doi:10.1038/ng.3970

[29] Abyzov A, Urban AE, Snyder M, Gerstein M. CNVnator: an approach to discover, genotype, and characterize typical and atypical CNVs from family and population genome sequencing. Genome Research. 2011;21(6):974–984. doi:10.1101/gr.114876.110

[30] Liao Y, Smyth GK, Shi W. The R package Rsubread is easier, faster, cheaper and better for alignment and quantification of RNA sequencing reads. Nucleic Acids Research. 2019;47(8):e47. doi:10.1093/nar/gkz114

[31] Magi A, Tattini L, Cifola I, D’Aurizio R, Benelli M, Mangano E, Battaglia C, Bonora E, Kurg A, Seri M, et al. EXCAVATOR: detecting copy number variants from whole-exome sequencing data. Genome Biology. 2013;14(10):R120. doi:10.1186/gb-2013-1410-r120

[32] Xiao F, Luo X, Hao N, Niu YS, Xiao X, Cai G, Amos CI, Zhang H. An accurate and powerful method for copy number variation detection. Bioinformatics. 2019;35(17):2891–2898. doi:10.1093/bioinformatics/bty1041

[33] Fromer M, Moran JL, Chambert K, Banks E, Bergen SE, Ruderfer DM, Handsaker RE, McCarroll SA, O’Donovan MC, Owen MJ, et al. Discovery and statistical genotyping of copy-number variation from whole-exome sequencing depth. American Journal of Human Genetics. 2012;91(4):597–607. doi:10.1016/j.ajhg.2012.08.005

[34] Neale BM, Kou Y, Liu L, Ma’ayan A, Samocha KE, Sabo A, Lin C-F, Stevens C, Wang L-S, Makarov V, et al. Patterns and rates of exonic de novo mutations in autism spectrum disorders. Nature. 2012;485(7397):242–245. doi:10.1038/nature11011

