## Supplementary Materials for "CN-RNN: a Deep Learning Framework for Copy Number Variation Detection with Exome Sequencing Data"

Corresponding to:

Feifei Xiao, Ph.D.

Department of Biostatistics, College of Public Health and Health Professions and College of Medicine, University of Florida

2004 Mowry Rd., CTRB building, Room 5227, Gainesville, FL, 32603

Outlines

### Supplementary Notes

#### Sequencing data preprocessing and normalization

Basic quality control steps were used with raw read counts, including removing exons with extreme GC content (<20% and >80%) or those with low mappability (<0.9). Then, a median-based normalization approach from the EXCAVATOR CNV detection method (Magi et al., 2013) was further applied. First, the exon mean read count (EMRC) was calculated using raw read counts ($RC$) and exon length ($L$): $EMRC=\frac{RC}{L}$. Each EMRC was then sequentially corrected for the three major sources of technical bias: (1) GC content, (2) mappability, and (3) exon size. The normalized read counts were defined as $\tilde{EMRC}_{i}=EMRC_{i}\frac{m}{m_{e}}$ where $m$ was the global median across all exons and $mₑ$ was the median of exons sharing the same $e$ value (e.g., GC content, mappability, or exon size).

We then computed the ratio of the normalized read counts to the reference samples, the logarithm transformation of which represents the main signal intensities (i.e., *log2R*). In the ASC dataset, ten healthy offspring controls with no family history of autism were selected as reference samples. In the PCGC dataset, twelve reference samples were selected and verified to contain no CNVs in the genomic regions corresponding to the validated CNVs in the test samples. For the 1000 Genomes Project dataset, ten samples with the lowest Gini index in their read depth data were selected as reference samples. Normalized read counts signals from the selected reference samples were pooled to estimate a population-level mean, which served as the baseline for calculating *log2R* values.

We further applied a local median-based smoothing procedure to reduce noise and suppress extreme signal fluctuations (Xiao et al., 2019). For each exon in each sample, a local window centered on that exon was defined using 10 neighboring exons on both sides. Within each local window, the mean, standard deviation, and median of the *log2R* values were calculated. An exon was classified as an outlier if its *log2R* value deviated from the local mean by more than twice the local standard deviation. Outlying values were then replaced by the corresponding local medians. This procedure reduced the influence of isolated extreme observations while preserving the broader regional copy number signal.

For the normalized read depth $R^{(i)}$, deviation from zero typically indicated duplications or deletions. To preserve local genomic context, a buffer zone of 20 exons was added to both sides of each candidate region. In addition, sequences were then zero-padded on both sides so that all input $R^{\left( i \right)}$ had uniform length, regardless of the number of exons spanned by the CNV events, which ensured consistent dimensionality across events. Because the normalized read-depth signal in diploid regions was centered near zero, zero-padding approximated the expected signal of flanking normal regions and did not introduce systematic bias into the classification.

For the genomic metadata, continuous variables (e.g., CNV length in kb, exon number) were log-transformed and Z-score standardized to place all continuous features on a comparable scale. Chromosome IDs were encoded as categorical variables.

#### Parallel read counting and CNV calling

To minimize the runtime of the preprocessing pipeline, the two most time-consuming steps were parallelized across CPU cores. Per-exon read counting from BAM files was performed using the *Rsubread featureCounts* function (Liao et al., 2019), with the *nthreads* argument set to distribute exon-level counting across multiple cores. The subsequent preliminary CNV calling (Qin et al., 2021) was likewise parallelized: per-chromosome CNV profiling tasks were dispatched concurrently across cores rather than processed sequentially. Because both steps scale near-linearly with the number of available cores, this configuration substantially reduced preprocessing wall time. These optimizations allowed CN-RNN's end-to-end runtime to remain shorter than that of other deep learning callers (Table 1).

### Supplementary Figures

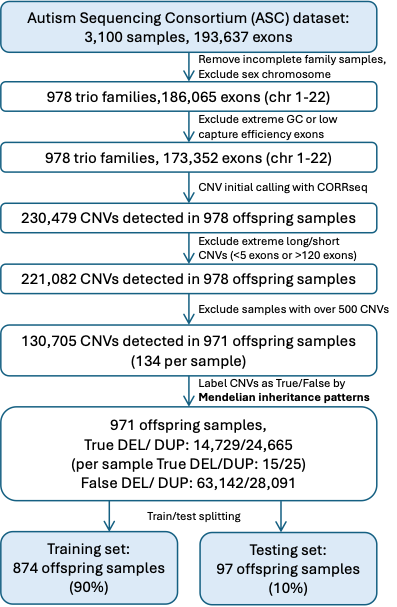

**Supplementary Figure 1. Overview of data processing and training label construction for the Autism Sequencing Consortium (ASC) dataset.** Whole-exome sequencing data from trio families were processed to generate CNV candidates, which were subsequently labeled as true or false CNVs based on the Mendelian rule of inheritance. These labeled CNVs were used to construct the training and internal validation datasets for CN-RNN. The figure outlines the study design of CNV calling including a detailed description of quality control steps, and summary of key numbers including the sample sizes and numbers of CNVs at each stage of analysis.

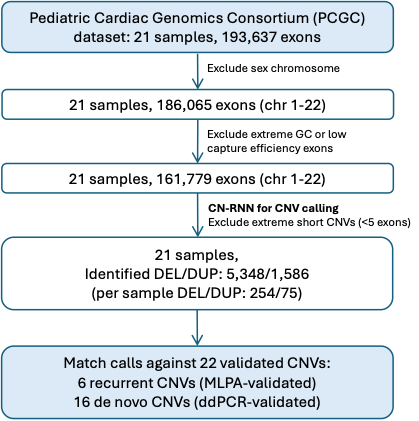

**Supplementary Figure 2. Overview of data processing for the Pediatric Cardiac Genomics Consortium (PCGC) dataset.** Whole-exome sequencing data were processed to generate CNV calls, which were evaluated against experimentally validated CNVs confirmed by MLPA or ddPCR. The figure outlines the CNV-calling study design, including a brief description of quality-control (QC) steps and a summary of key counts (sample size and number of CNVs at each stage of analysis).

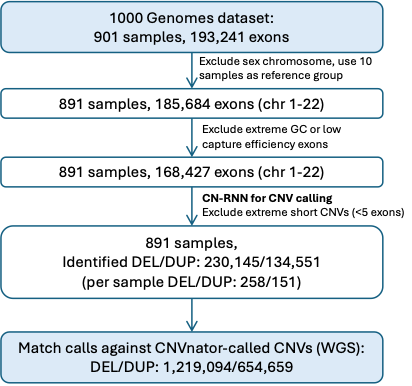

**Supplementary Figure 3. Overview of data processing for the 1000 Genomes Project dataset.** Whole-exome sequencing data were processed to generate CNV calls which were subsequently evaluated against WGS data generated CNV calls. The figure outlines the CNV-calling study design, including a brief description of quality-control (QC) steps and a summary of key counts (sample size and number of CNVs at each stage of analysis).

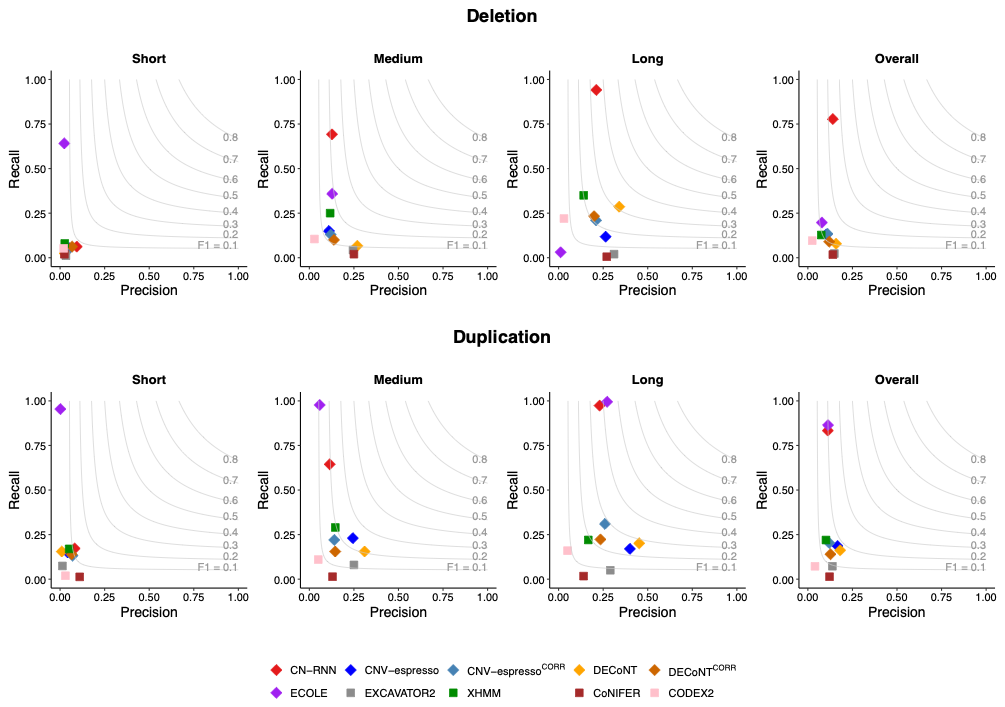

**Supplementary Figure 4. Length-stratified performance on the 1000 Genomes Project dataset.** Precision-recall scatter plots of CN-RNN and competing methods on the 1000 Genomes Project WES dataset, stratified by CNV length: short (5–10 exons), medium (11–40 exons), long (>40 exons), and overall (all lengths pooled). Results are shown for deletions (top row) and duplications (bottom row). CNV-espresso and DECoNT (without superscript) used XHMM as the initial CNV caller. CNV-espresso^CORR^ and DECoNT^CORR^ used CORRseq as the initial CNV caller.

### Supplementary Tables

**Supplementary Table 1. Summary of the CNV training set** **in the ASC trio dataset.** The table summarizes the training set by CNV type (deletions and duplications), training label (true CNVs, false CNVs, and negative controls), and CNV length (number of exons and length in kilobase[kb]). Training labels were assigned at a designed ratio of 50:40:10 (true CNVs: false CNVs: negative controls) to maintain a balanced training set. Del: Deletions. Dup: Duplications; NC: negative controls.

| **CNV type** | **Training label** | **Stratified by length (in kb)** | | | **Stratified by number of exons** | | | **Total count** |
| --- | --- | --- | --- | --- | --- | --- | --- | --- |
|  |  | **Short**  **(0-50 kb)** | **Medium**  **(50-200 kb)** | **Long (200-5,000 kb)** | **Short**  **(5-10)** | **Medium**  **(11-40)** | **Long**  **(41-120)** |  |
| Del | True | 4,382 | 4,261 | 4,357 | 4,612 | 6,138 | 2,250 | **13,000** |
|  | False | 4,485 | 3,273 | 2,642 | 4,959 | 4,644 | 797 | **10,400** |
|  | NC | 1,195 | 717 | 688 | 1,151 | 1,184 | 265 | **2,600** |
| Dup | True | 9,335 | 6,898 | 5,767 | 8,708 | 10,394 | 2,898 | **22,000** |
|  | False | 7,946 | 5,090 | 4,564 | 7,241 | 8,461 | 1,898 | **17,600** |
|  | NC | 1,929 | 1,182 | 1,289 | 1,754 | 2,095 | 551 | **4,400** |

**Supplementary Table 2. Summary of the CN-RNN^EXCAVATOR2^ training set derived from EXCAVATOR2 CNV calls in the ASC trio dataset.** The table summarizes the training set by CNV type (deletions and duplications), training label (true CNVs, false CNVs, and negative controls), and CNV length (number of exons and length in kilobases [kb]). This training set was used to train CN-RNN^EXCAVATOR2^, a variant model sharing the same hybrid neural network architecture as CN-RNN. Training labels were assigned at the same designed ratio of 50:40:10 (true CNVs: false CNVs: negative controls) to maintain a balanced training set. Del: Deletions. Dup: Duplications.

| **CNV type** | **Training label (proportion)** | **Stratified by length (in kb)** | | | **Stratified by number of exons** | | | **Total count** |
| --- | --- | --- | --- | --- | --- | --- | --- | --- |
|  |  | **Short**  **(0-50 kb)** | **Medium**  **(50-200 kb)** | **Long (200-5,000 kb)** | **Short**  **(5-10)** | **Medium**  **(11-40)** | **Long**  **(41-120)** |  |
| Del | True CNV (50%) | 2,434 | 4,512 | 3,053 | 3,840 | 4,684 | 1,476 | **10,000** |
|  | False CNV (40%) | 1,987 | 2,872 | 3,141 | 2,201 | 4,000 | 1,799 | **8,000** |
|  | Negative control (10%) | 482 | 892 | 626 | 768 | 937 | 295 | **2,000** |
| Dup | True CNV (50%) | 316 | 2,244 | 3,439 | 1,033 | 1,930 | 3,037 | **6,000** |
|  | False CNV (40%) | 345 | 1,657 | 2,798 | 1,044 | 1,435 | 2,321 | **4,800** |
|  | Negative control (10%) | 64 | 436 | 700 | 206 | 386 | 408 | **1,200** |

**Supplementary Table 3. Summary of genomic metadata features used as input to the CN-RNN model for CNV classification.**

| **Feature name** | **Range** | **Description** |
| --- | --- | --- |
| Number_Exon | [5, 120] | Number of exons |
| Length | [1.20, 5,000] | Physical length in kilobases (kb) |
| median_GC | [0.267, 0.765] | Median GC content across the CNV region |
| median_mappability | [0.920, 1.000] | Median mappability across the CNV region |
| L2R_mean | [-5.676, 1.636] | Mean log2 ratio of normalized coverage signal |
| L2R_sd | [0.019, 3.435] | Standard deviation of log2 ratio within the CNV |
| Chr_id | {1,…, 22} | Chromosome ids |

**Supplementary Table 4. Performance comparison on the ASC trio dataset, stratified by CNV size (number of exons).** Metrics include precision, recall, and F1 score and correspond to the results shown in Figure 2.

| **Model** | **Length**  **(number of exons)** | **Deletion** | | | **Duplication** | | |
| --- | --- | --- | --- | --- | --- | --- | --- |
|  |  | **F1-score** | **Precision** | **Recall** | **F1-score** | **Precision** | **Recall** |
| **CN-RNN** | Short (5-10) | **0.65** | 0.68 | 0.63 | **0.72** | 0.72 | 0.73 |
|  | Medium (11-40) | **0.72** | 0.75 | 0.70 | **0.72** | 0.71 | 0.74 |
|  | Long (> 41) | **0.83** | 0.75 | 0.94 | **0.79** | 0.74 | 0.84 |
|  | Overall | **0.71** | 0.72 | 0.71 | **0.73** | 0.72 | 0.75 |
| **CN-RNN^EXCAVATOR2^** | Short (5-10) | **0.61** | 0.63 | 0.60 | **0.66** | 0.63 | 0.68 |
|  | Medium (11-40) | **0.66** | 0.67 | 0.65 | **0.69** | 0.68 | 0.70 |
|  | Long (> 41) | **0.68** | 0.69 | 0.68 | **0.72** | 0.71 | 0.72 |
|  | Overall | **0.65** | 0.65 | 0.64 | **0.68** | 0.66 | 0.70 |
| **CNV-espresso** | Short (5-10) | **0.27** | 0.41 | 0.20 | **0.32** | 0.30 | 0.34 |
|  | Medium (11-40) | **0.26** | 0.27 | 0.25 | **0.44** | 0.31 | 0.74 |
|  | Long (> 41) | **0.16** | 0.26 | 0.12 | **0.59** | 0.46 | 0.84 |
|  | Overall | **0.25** | 0.32 | 0.21 | **0.42** | 0.33 | 0.60 |
| **DECoNT** | Short (5-10) | **0.23** | 0.22 | 0.23 | **0.19** | 0.28 | 0.14 |
|  | Medium (11-40) | **0.33** | 0.27 | 0.42 | **0.21** | 0.31 | 0.16 |
|  | Long (> 41) | **0.49** | 0.34 | 0.89 | **0.61** | 0.45 | 0.92 |
|  | Overall | **0.32** | 0.26 | 0.42 | **0.29** | 0.32 | 0.26 |
| **ECOLE** | Short (5-10) | **0.19** | 0.17 | 0.22 | **0.08** | 0.06 | 0.16 |
|  | Medium (11-40) | **0.28** | 0.23 | 0.36 | **0.20** | 0.12 | 0.68 |
|  | Long (> 41) | **0.55** | 0.43 | 0.78 | **0.30** | 0.18 | 0.87 |
|  | Overall | **0.28** | 0.23 | 0.36 | **0.17** | 0.10 | 0.50 |
| **CORRseq** | Short (5-10) | **0.37** | 0.23 | 1.00 | **0.44** | 0.28 | 1.00 |
|  | Medium (11-40) | **0.43** | 0.28 | 1.00 | **0.46** | 0.30 | 1.00 |
|  | Long (> 41) | **0.52** | 0.35 | 1.00 | **0.59** | 0.42 | 1.00 |
|  | Overall | **0.42** | 0.27 | 1.00 | **0.47** | 0.31 | 1.00 |
| **EXCAVATOR2** | Short (5-10) | **0.35** | 0.21 | 1.00 | **0.23** | 0.13 | 1.00 |
|  | Medium (11-40) | **0.39** | 0.25 | 1.00 | **0.25** | 0.14 | 1.00 |
|  | Long (> 41) | **0.47** | 0.31 | 1.00 | **0.25** | 0.15 | 1.00 |
|  | Overall | **0.39** | 0.24 | 1.00 | **0.24** | 0.14 | 1.00 |
| **XHMM** | Short (5-10) | **0.13** | 0.39 | 0.08 | **0.24** | 0.43 | 0.17 |
|  | Medium (11-40) | **0.16** | 0.39 | 0.10 | **0.27** | 0.45 | 0.19 |
|  | Long (> 41) | **0.19** | 0.44 | 0.12 | **0.30** | 0.49 | 0.22 |
|  | Overall | **0.15** | 0.40 | 0.10 | **0.26** | 0.45 | 0.19 |
| **CoNIFER** | Short (5-10) | **0.02** | 0.32 | 0.01 | **0.02** | 0.11 | 0.01 |
|  | Medium (11-40) | **0.04** | 0.38 | 0.02 | **0.02** | 0.13 | 0.01 |
|  | Long (> 41) | **0.04** | 0.39 | 0.02 | **0.04** | 0.14 | 0.02 |
|  | Overall | **0.03** | 0.36 | 0.02 | **0.02** | 0.12 | 0.01 |
| **CODEX2** | Short (5-10) | **0.04** | 0.02 | 0.16 | **0.05** | 0.03 | 0.09 |
|  | Medium (11-40) | **0.05** | 0.03 | 0.19 | **0.07** | 0.05 | 0.11 |
|  | Long (> 41) | **0.05** | 0.03 | 0.22 | **0.08** | 0.05 | 0.16 |
|  | Overall | **0.05** | 0.03 | 0.18 | **0.06** | 0.04 | 0.11 |

**Supplementary Table 5. CNV validation results in the PCGC dataset using CN-RNN.** Performance was evaluated against experimentally validated CNVs confirmed by MLPA or ddPCR. CNV Type: true CNV type.

| **Blinded ID** | **Chromosome** | **Location** | **CNV type** | **CN-RNN prediction** | **Size (kb)** | **CNV Category** | **Reference** |
| --- | --- | --- | --- | --- | --- | --- | --- |
| 1-03171 | 1 | 145586403-145799634 | Dup | Dup | 213.20 | Rare | Glessner et al., 2014. |
| 1-01036 | 1 | 146631133-147416212 | Dup | Dup | 785.10 | Rare | Glessner et al., 2014. |
| 1-01401 | 2 | 102493466-103001458 | Del | Del | 508.00 | Rare | Glessner et al., 2014. |
| 1-01401 | 2 | 145155868-145274931 | Del | Del | 119.10 | Rare | Glessner et al., 2014. |
| 1-00771 | 4 | 185603346-185638397 | Del | Del | 35.10 | Rare | Glessner et al., 2014. |
| 1-00977 | 7 | 138258252-143807632 | Del | Del | 5549.30 | Rare | Glessner et al., 2014. |
| 1-00566 | 8 | 11606428-11710963 | Del | Non-CNV | 104.50 | Rare | Glessner et al., 2014. |
| 1-00230 | 11 | 86939592-87025456 | Del | Del | 85.90 | Rare | Glessner et al., 2014. |
| 1-01486 | 11 | 125641368-134943190 | Del | Del | 9301.80 | Rare | Glessner et al., 2014. |
| 1-01396 | 15 | 22750305-23228712 | Del | Del | 478.40 | Rare | Glessner et al., 2014. |
| 1-00243 | 15 | 22835893-23062345 | Del | Del | 226.50 | Rare | Glessner et al., 2014. |
| 1-01994 | 15 | 28389771-28446734 | Del | Non-CNV | 57.00 | Rare | Glessner et al., 2014. |
| 1-01696 | 15 | 44833588-44856873 | Del | Del | 23.30 | Rare | Glessner et al., 2014. |
| 1-00113 | 22 | 18886915-22000000 | Del | Del | 3113.10 | Rare | Glessner et al., 2014. |
| 1-01836 | 22 | 19020529-21380382 | Del | Non-CNV | 2359.90 | Rare | Glessner et al., 2014. |
| 1-00425 | 22 | 36038076-36149338 | Del | Del | 111.30 | Rare | Glessner et al., 2014. |
| CG0003-3939 | 7 | 7q11.23 | Del | Del | - | Recurrent | dbGaP phs000571.v7.p3 |
| CG0004-1809 | 22 | 22q11.2 | Dup | Non-CNV | - | Recurrent | dbGaP phs000571.v7.p3 |
| CG0023-6046 | 22 | 22q11.2 | Del | Del | - | Recurrent | dbGaP phs000571.v7.p3 |
| CG0019-6438 | 22 | 22q11.2 | Dup | Dup | - | Recurrent | dbGaP phs000571.v7.p3 |
| CG0014-5582 | 22 | 22q11.2 | Del | Del | - | Recurrent | dbGaP phs000571.v7.p3 |
| CG0011-0164 | 22 | 22q11.2 | Del | Non-CNV | - | Recurrent | dbGaP phs000571.v7.p3 |

**Supplementary Table 6. Performance evaluation in the 1000 Genomes Project dataset.** WES-based CNV predictions were evaluated against CNVnator-derived WGS CNVs, corresponding to the results shown in Figure 4. CNV-espresso: CNV-espresso applied to XHMM calls. CNV-espresso^CORR^: CNV-espresso applied to CORRseq calls. DECoNT: DECoNT applied to XHMM calls. DECoNT^CORR^: DECoNT applied to CORRseq calls.

| **Model** | **Length**  **(number of exons)** | **Deletion** | | | **Duplication** | | |
| --- | --- | --- | --- | --- | --- | --- | --- |
|  |  | **F1-score** | **Precision** | **Recall** | **F1-score** | **Precision** | **Recall** |
| **CN-RNN** | Short (5-10) | **0.07** | 0.09 | 0.06 | **0.11** | 0.08 | 0.17 |
|  | Medium (11-40) | **0.21** | 0.13 | 0.69 | **0.19** | 0.11 | 0.64 |
|  | Long (> 41) | **0.35** | 0.21 | 0.94 | **0.37** | 0.23 | 0.97 |
|  | Overall | **0.24** | 0.14 | 0.78 | **0.20** | 0.11 | 0.83 |
| **CNV-espresso** | Short (5-10) | **0.05** | 0.05 | 0.05 | **0.07** | 0.04 | 0.15 |
|  | Medium (11-40) | **0.13** | 0.11 | 0.15 | **0.24** | 0.24 | 0.23 |
|  | Long (> 41) | **0.16** | 0.26 | 0.12 | **0.24** | 0.40 | 0.17 |
|  | Overall | **0.12** | 0.10 | 0.13 | **0.18** | 0.17 | 0.19 |
| **CNV-espresso^CORR^** | Short (5-10) | **0.04** | 0.04 | 0.04 | **0.09** | 0.07 | 0.13 |
|  | Medium (11-40) | **0.12** | 0.12 | 0.13 | **0.17** | 0.14 | 0.22 |
|  | Long (> 41) | **0.21** | 0.21 | 0.21 | **0.28** | 0.26 | 0.31 |
|  | Overall | **0.12** | 0.11 | 0.13 | **0.15** | 0.12 | 0.20 |
| **DECoNT** | Short (5-10) | **0.03** | 0.03 | 0.02 | **0.02** | 0.01 | 0.16 |
|  | Medium (11-40) | **0.11** | 0.27 | 0.07 | **0.21** | 0.31 | 0.16 |
|  | Long (> 41) | **0.31** | 0.34 | 0.29 | **0.28** | 0.45 | 0.20 |
|  | Overall | **0.10** | 0.16 | 0.08 | **0.17** | 0.18 | 0.16 |
| **DECoNT^CORR^** | Short (5-10) | **0.06** | 0.07 | 0.06 | **0.09** | 0.07 | 0.14 |
|  | Medium (11-40) | **0.12** | 0.14 | 0.10 | **0.15** | 0.15 | 0.16 |
|  | Long (> 41) | **0.21** | 0.20 | 0.23 | **0.23** | 0.23 | 0.22 |
|  | Overall | **0.10** | 0.12 | 0.09 | **0.13** | 0.13 | 0.14 |
| **ECOLE** | Short (5-10) | **0.05** | 0.02 | 0.64 | **0.00** | 0.00 | 0.95 |
|  | Medium (11-40) | **0.19** | 0.13 | 0.36 | **0.11** | 0.06 | 0.98 |
|  | Long (> 41) | **0.02** | 0.01 | 0.03 | **0.43** | 0.27 | 0.99 |
|  | Overall | **0.11** | 0.08 | 0.20 | **0.20** | 0.11 | 0.86 |
| **EXCAVATOR2** | Short (5-10) | **0.02** | 0.03 | 0.01 | **0.02** | 0.01 | 0.07 |
|  | Medium (11-40) | **0.07** | 0.25 | 0.04 | **0.12** | 0.25 | 0.08 |
|  | Long (> 41) | **0.04** | 0.31 | 0.02 | **0.09** | 0.29 | 0.05 |
|  | Overall | **0.04** | 0.15 | 0.02 | **0.10** | 0.14 | 0.07 |
| **XHMM** | Short (5-10) | **0.04** | 0.03 | 0.08 | **0.08** | 0.05 | 0.17 |
|  | Medium (11-40) | **0.16** | 0.12 | 0.25 | **0.19** | 0.15 | 0.29 |
|  | Long (> 41) | **0.20** | 0.14 | 0.35 | **0.19** | 0.17 | 0.22 |
|  | Overall | **0.09** | 0.08 | 0.13 | **0.14** | 0.10 | 0.22 |
| **CoNIFER** | Short (5-10) | **0.02** | 0.02 | 0.02 | **0.02** | 0.11 | 0.01 |
|  | Medium (11-40) | **0.04** | 0.25 | 0.02 | **0.03** | 0.13 | 0.01 |
|  | Long (> 41) | **0.01** | 0.27 | 0.01 | **0.03** | 0.14 | 0.02 |
|  | Overall | **0.03** | 0.14 | 0.02 | **0.03** | 0.12 | 0.01 |
| **CODEX2** | Short (5-10) | **0.03** | 0.02 | 0.05 | **0.02** | 0.03 | 0.02 |
|  | Medium (11-40) | **0.04** | 0.03 | 0.11 | **0.07** | 0.05 | 0.11 |
|  | Long (> 41) | **0.05** | 0.03 | 0.22 | **0.08** | 0.05 | 0.16 |
|  | Overall | **0.04** | 0.02 | 0.10 | **0.05** | 0.04 | 0.07 |
